# Frontotemporal Dementia Patient Neurons With Progranulin Deficiency Display Protein Dyshomeostasis

**DOI:** 10.1101/2023.01.18.524611

**Authors:** Lisa Elia, Bianca Herting, Amela Alijagic, Christina Buselli, Leela Wong, Grace Morrison, Miguel A. Prado, Joao A. Paulo, Steven P. Gygi, Daniel Finley, Steven Finkbeiner

**Author notes:** **Corresponding authors:** Lisa P. Elia and Steven Finkbeiner, Gladstone Institutes, 1650 Owens St., San Francisco, California 94158.

## Abstract

Haploinsufficiency of progranulin (PGRN) causes frontotemporal dementia (FTD), a devastating neurodegenerative disease with no effective treatment. PGRN is required for efficient proteostasis, as loss of neuronal PGRN results in dysfunctional lysosomes and impaired clearance and cytoplasmic aggregation of TDP-43, a protein involved in neurodegeneration in FTD. These and other events lead to neurodegeneration and neuroinflammation. However, the detailed mechanisms leading to protein dyshomeostasis in PGRN-deficient cells remain unclear. We report here the development of human cell models of FTD with PGRN-deficiency to explore the molecular mechanisms underlying proteostasis breakdown and TDP-43 aggregation in FTD. Neurons differentiated from FTD patient induced pluripotent stem cells (iPSCs) have reduced PGRN levels, and the neurons recapitulate key disease features, including impaired lysosomal function, defective TDP-43 turnover and accumulation, neurodegeneration, and death. Proteomic analysis revealed altered levels of proteins linked to the autophagy-lysosome pathway (ALP) and the ubiquitin-proteasome system (UPS) in FTD patient neurons, providing new mechanistic insights into the link between PGRN-deficiency and disease pathobiology.

## Introduction

Haploinsufficiency in the *GRN* gene causes the fatal neurodegenerative disease frontotemporal dementia (FTD), an early-onset, presenile dementia that typically presents before the age of 65, and is characterized by language impairment and behavioral changes. Over 60 mutations in the *GRN* locus result in functional null alleles that lead to haploinsufficient levels of the encoded protein, progranulin (PGRN). PGRN is a secreted growth factor with diverse functions, including regulating the immune system and brain function. Dysfunction of PGRN has been linked to cancer, metabolic disorders, and multiple neurodegenerative diseases in addition to FTD. Decreased serum levels of PGRN caused by a single nucleotide polymorphism (SNP) in the 3’-UTR of *GRN* confers an elevated risk of developing Alzheimer’s disease (AD), and increasing the levels of PGRN in mouse models of AD suppresses neuroinflammatory-, pathological-, and cognitive disease phenotypes^1-8^. A risk factor SNP for Parkinson’s disease (PD) has been identified in the *GRN* locus based on genome-wide association studies (GWAS) ^9^. A novel mutation in *GRN* was shown to promote not only TDP-43, but also tau and α-synuclein aggregation pathology in two families presenting with variable clinical features consistent with FTD and parkinsonism^10^.

Nullizygous mutations in *GRN* lead to the development of neuronal ceroid lipofuscinosis (NCL), a lysosomal storage disorder (LSD)^2, 11-14^. PGRN is also implicated in Gaucher’s disease, a common lysosomal storage disease caused by mutations in *GBA1*, which encodes the lysosomal enzyme glucocerebrosidase (GCase)^15^. PGRN has been shown recently to act as a co-chaperone, along with HSP70, for GCase^16^, promoting its biosynthetic delivery to the lysosome.

To investigate the link between PGRN-deficiency to lysosome dysfunction, protein dyshomeostasis and neurodegeneration, we and others have employed cultured murine *GRN*-deficient cortical neuronal models of FTD. We found that *GRN*-deficient cortical neurons display deficits in protein homeostasis (proteostasis) and disruption of the autophagy lysosome pathway(s) (ALP)^17^. We reported that PGRN-deficient lysosomes accumulated undigested material, including lipofuscin, and became enlarged indicating impaired lysosomal function. Using a photoswitchable autophagy flux bioreporter, we found, paradoxically, that PGRN-deficient cortical neurons display enhanced basal autophagic flux^17^. We proposed that the accelerated autophagic clearance may be a compensatory mechanism responding to impaired lysosomal function.

Reduced levels of PGRN in the brain lead to excessive neuroinflammation with an increased presence of both activated microglia (microgliosis) and reactive astrocytes (astrogliosis). Neurodegeneration and neuronal death occur predominantly within the frontal and temporal cortices of PGRN-deficient FTD patients. Brain tissue from FTD patients with *GRN* haploinsufficiency also shows increased cytoplasmic accumulation of ubiquitinated proteins in aggregates, together with TDP-43, which is normally localized to the nucleus. Cytoplasmic mislocalization of TDP-43 has been linked to neurodegeneration in both FTD and another neurodegenerative disease, ALS.

We have previously reported that cultured murine *GRN*-deficient cortical neurons recapitulated the increased cytoplasmic accumulation of TDP-43. Misfolded TDP-43 is cleared by the ALP^23^, but others have reported that both chaperone-mediated autophagy (CMA) and the ubiquitin-proteasome system (UPS) may also clear physiological and pathogenic forms of TDP-43^18-21^. A better understanding of the molecular mechanisms contributing to the breakdown of proteostasis and accumulation of toxic TDP-43 in neurons may enable the development of therapeutic strategies to treat FTD.

Mouse models of *Grn*-haploinsufficient FTD have been used to better understand the biological functions of PGRN and pathological changes associated with PGRN-deficiency^22-28^. They also provide preclinical tools to test novel therapeutics. However, animal models lack the genetic background of human FTD patients, and the underlying genetics of humans may be critical in understanding the cellular mechanisms of how PGRN deficiency leads to behavioral dysfunction, neurodegeneration and death of FTD patients.

We report here the development of a human PGRN-deficient FTD neuronal model consisting of induced pluripotent stem cells (iPSCs)-derived neurons (i-neurons) from a patient carrying a common FTD-linked mutation in the *GRN* locus (*GRN(R493X)*). The patient-derived i-neurons had reduced PGRN levels and recapitulated key FTD disease features, including enlarged lysosomes, defects in lysosomal acidification, and deficits in both TDP-43 and α-synuclein clearance. In addition, the i-neurons exhibit neurite loss, and neuronal death. Quantitative tandem mass tagging (TMT)-proteomics analysis of PGRN-deficient FTD patient i-neurons revealed aberrant changes in the levels of proteins linked to both the autophagy-lysosome (ALP) and the ubiquitin-proteasome system (UPS) clearance pathways. In particular, PGRN-deficient FTD patient i-neurons exhibited altered levels of ATP6V0A2 and ATP6V1D, two subunits of the v-ATPase that are responsible for efficient lysosome acidification; altered levels of the lysosomal enzymes HEXA and HEXB, which are important for ganglioside degradation; and altered levels of the lysosomal cathepsins C and Z. We also found changes in proteins linked to the UPS pathway, including three proteasome activators:, PSME1, PSME2, and ZFAND1. These data suggest that PGRN-deficiency may result in the impairment of multiple proteostasis pathways leading, in turn, to aggregation of neurodegenerative disease proteins to contribute to disease pathogenesis.

## Results

### FTD i-neurons spontaneously degenerate over time

To create human FTD cell models, we differentiated iPSCs into neural precursors cells (NPCs) and then forebrain cortical neurons from a patient harboring the most common *GRN* mutation that causes FTD, the *GRN(R493X)* mutation (FTD(*GRN(R493X)*). We also generated i-neurons from a healthy control individual (WTC11) using a dual SMAD inhibition monolayer approach. The PGRN-deficient FTD patient i-neurons had a significant reduction in the levels of secreted PGRN measured by ELISA assay (**Fig.1A**) consistent with PGRN-deficiency observed in the disease.

Neuronal morphology of the PGRN-deficient FTD i-neurons (**Fig. 1**) showed dystrophic neurites with sparse, short, and thickened features similar to neurons described in the brains of PGRN-deficient FTD patients^29^. We found that the PGRN-deficient FTD i-neurons had overly simplified neurite arbors (**Fig. 1B, C**) with a significant reduction in neurite length (**Fig. 1D**), suggesting that the PGRN-deficient FTD i-neurons may undergo neurodegeneration.

**Figure 1.**
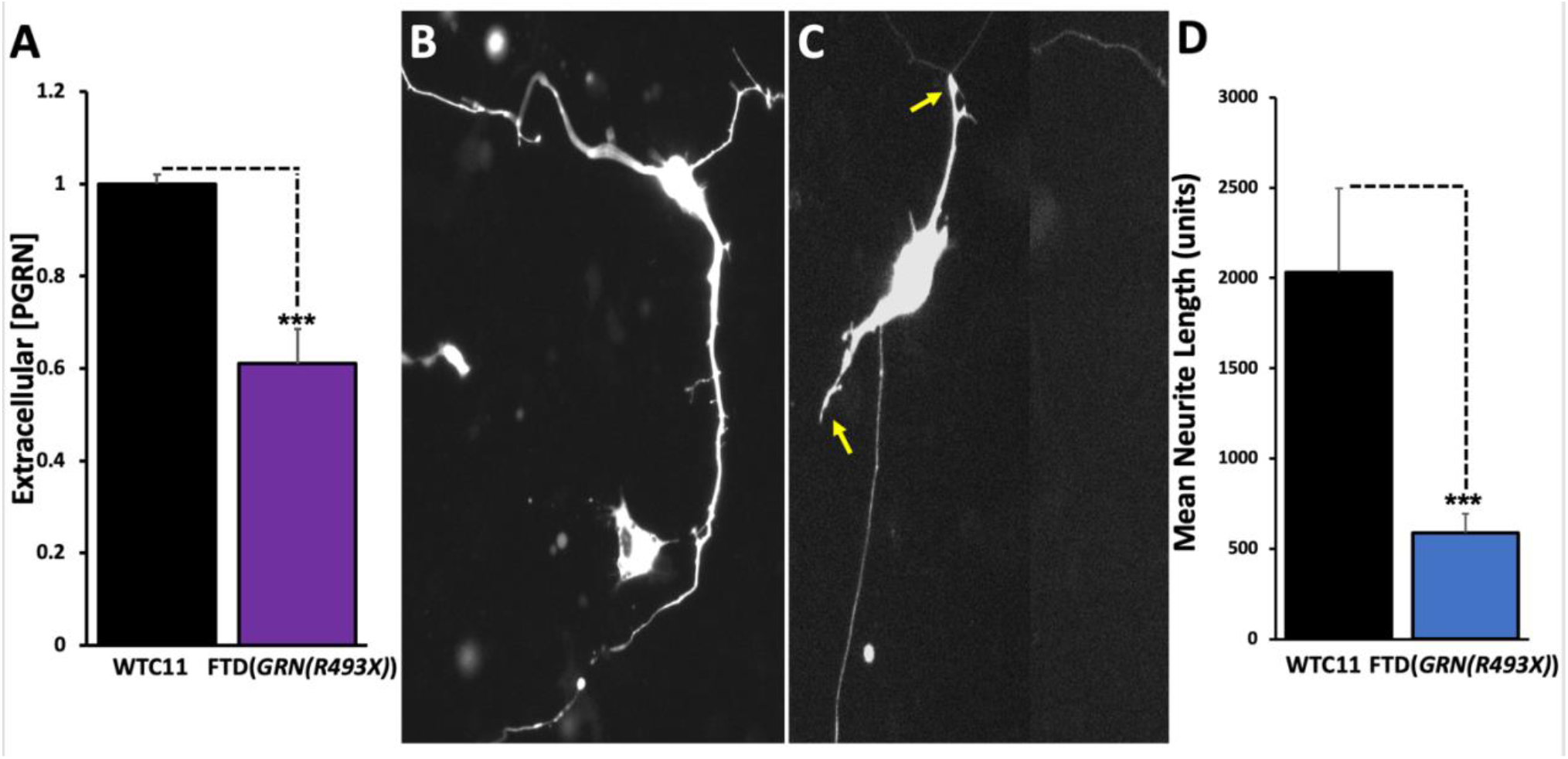
Reduced PGRN leads to neurite degeneration in human FTD patient iPSC-differentiated i-neurons. **A**. Quantitative ELISA of extracellular (secreted) PGRN in human healthy control (WTC11) or FTD patient (FTD(*GRN(R493X)*) i-neurons. **B, C**. i-Neurons cultured from a human FTD patient iPSC line (FTD(*GRN(R493X)*) or a healthy control (WTC11) line were transfected with a plasmid expressing EGFP under the human synapsin1 promoter (pGW1-hSyn1-EGFP) to visualize overall neuronal morphology. Yellow arrows (**C**) indicate degenerating dendrites in the FTD patient i-neurons. **D**. Quantitation of mean neurite length per cell. Statistical tests: Student’s t-test (***p<0.001).

The PGRN-deficient FTD i-neurons degenerated and died prematurely (**Fig. 2**). We used robotic microscopy (RM), a unique single cell longitudinal automated fluorescence microscopy platform that we developed, to determine the risk of death for the PGRN-deficient FTD i-neurons^30-32^. RM can be used to measure the lifespan of individual neurons expressing a fluorescent cell morphology marker, such as EGFP driven by the synapsin1 promoter. Cell death is indicated by the loss of EGFP fluorescence due to the disruption of cell integrity. We calculated the cumulative risk of death for each population of i-neurons and found that PGRN-deficiency increased the risk of death by ∼257%, suggesting that survival of PGRN-deficient FTD i-neurons was significantly impaired (**Fig. 2**).

**Figure 2.**
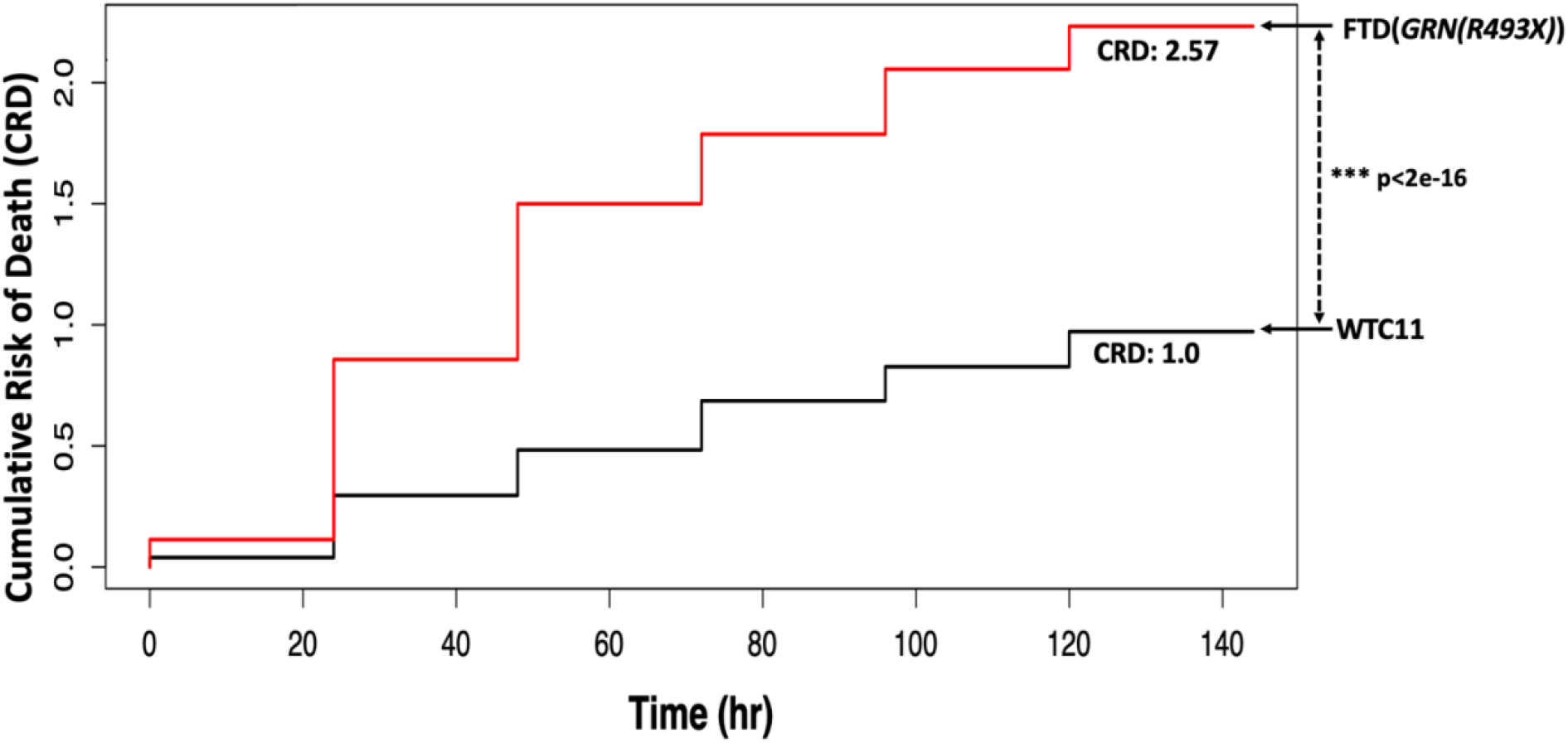
PGRN-deficient FTD patient i-neurons display increased neurotoxicity. Kaplan-Meier survival analysis was used to determine the cumulative risk of death functions for each population of transduced i-neurons. The y-axis represents a quantitative measure of the accumulated risk of cell death overtime. ***p <2e^-16^ (log-rank test). The cumulative risk of death represents the probability that cells in a cohort will die over the time interval for each experiment. Cumulative risk of death curves compiled from three separate experiments, including a total of 38,382 FTD(*GRN(R493X)*) patient i-neurons or 36,106 healthy control WTC11 i-neurons expressing EGFP under the human synapsin1 promoter.

### PGRN-deficient FTD i-neurons do not efficiently clear TDP-43

PGRN is important for maintaining the functional integrity of the normally nuclear localized protein, TDP-43. Cytoplasmic mislocalized and aggregated TDP-43 results in significant cellular toxicity either through a loss-of-function of its essential nuclear role in regulating RNA processing or a gain-of-function toxic event in the cytoplasm, or both. TDP-43 mislocalization is a key cytopathological feature of PGRN-deficient FTD. We previously reported that reduced PGRN levels resulted in TDP-43 cytoplasmic mislocalization and aggregation in cultured rodent PGRN-deficient FTD model neurons. Similarly, we examined whether the human PGRN-deficient FTD i-neurons exhibit TDP-43 pathology using immunocytochemistry and found that in the healthy control i-neurons, TDP-43 was restricted to the nucleus (**Fig. 3 A-G, O**) whereas the PGRN-deficient FTD i-neurons had a significant increase in TDP-43 accumulation in the cytoplasm (**Fig. 3 H-N, O**).

**Figure 3:**
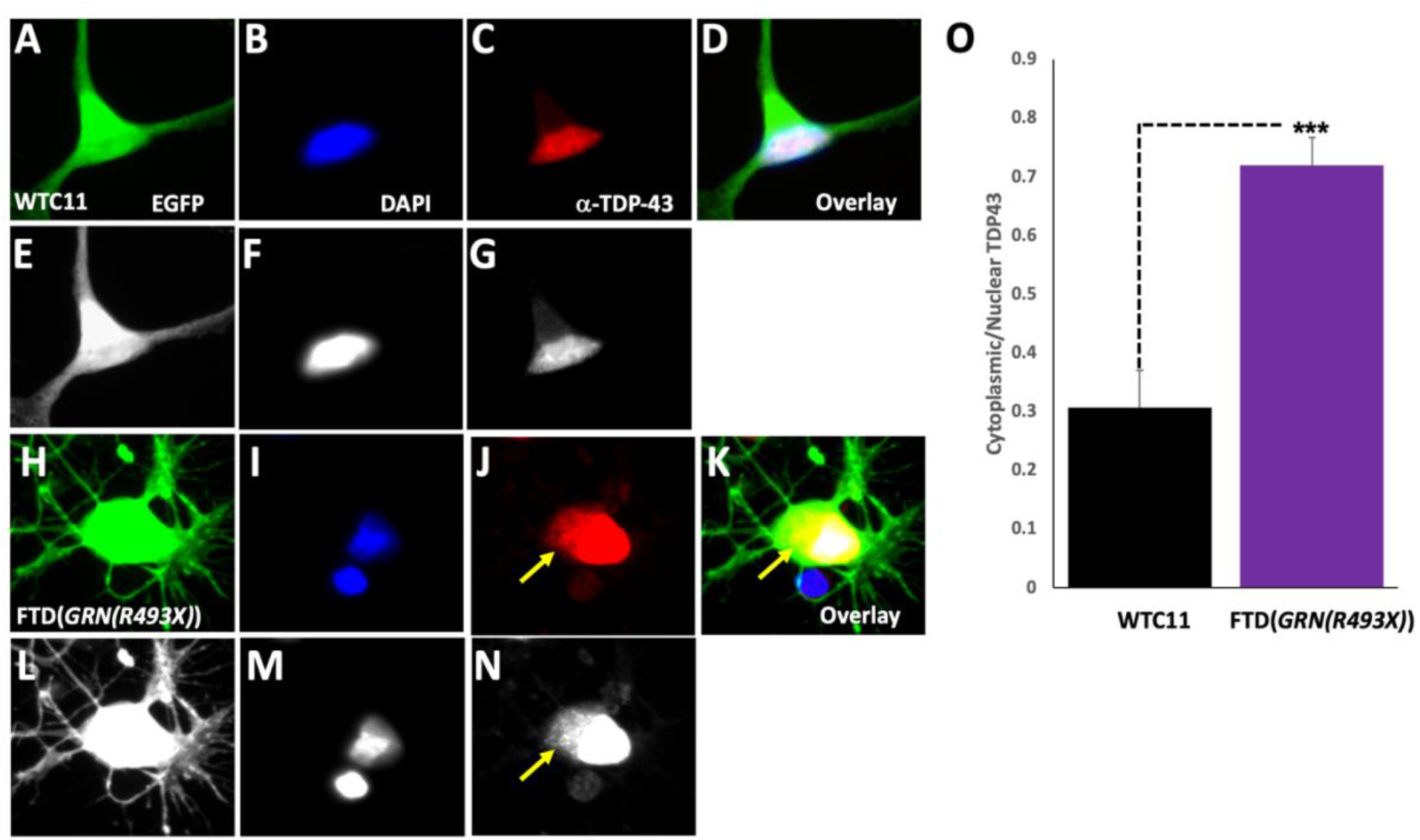
PGRN-deficient FTD patient cortical i-neurons exhibit cytoplasmic TDP-43 accumulation. Images of cultured human cortical i-neurons transduced with a lentivirus expressing EGFP (green) under the human synapsin1 promoter fixed and stained with DAPI (blue) and a TDP-43 antibody (red); corresponding grayscale images are also shown. **(A-G)**: representative images of healthy control (WTC11) cortical i-neurons. **(H-N)**: representative images of FTD patient (*GRN(R493X)*) cortical i-neurons. Yellow arrows indicate cytoplasmic accumulation of TDP-43 in the FTD patient i-neurons (**J, K, N**). **(O)** Quantitation of the ratio of cytoplasmic:nuclear TDP-43. Statistical test: Student’s t-test. (***p<0.001).

We next used a sensitive optical pulse labeling (OPL) method and longitudinal imaging with RM to quantify the rate of TDP-43 turnover at a single cell level in the PGRN-deficient FTD i-neurons (**Fig. 4**)^31^. We used a photoswitchable TDP-43 reporter, TDP-43-Dendra2, which allows us to irreversibly photoconvert a population of intracellular TDP-43 protein and then subsequently tracked the rate of its clearance in single cells, and calculate the average half-life of TDP-43 in this uniquely tagged population. Healthy control i-neurons had an average TDP-43-Dendra2 half-life of ∼29h, whereas the PGRN-deficient FTD i-neurons had an average TDP-43-Dendra2 half-life of ∼66h. This indicates that PGRN-deficient FTD i-neurons were significantly less efficient at clearing TDP-43-Dendra2 than healthy control i-neurons, consistent with the increased accumulation of TDP-43 in the disease i-neurons.

**Figure 4:**
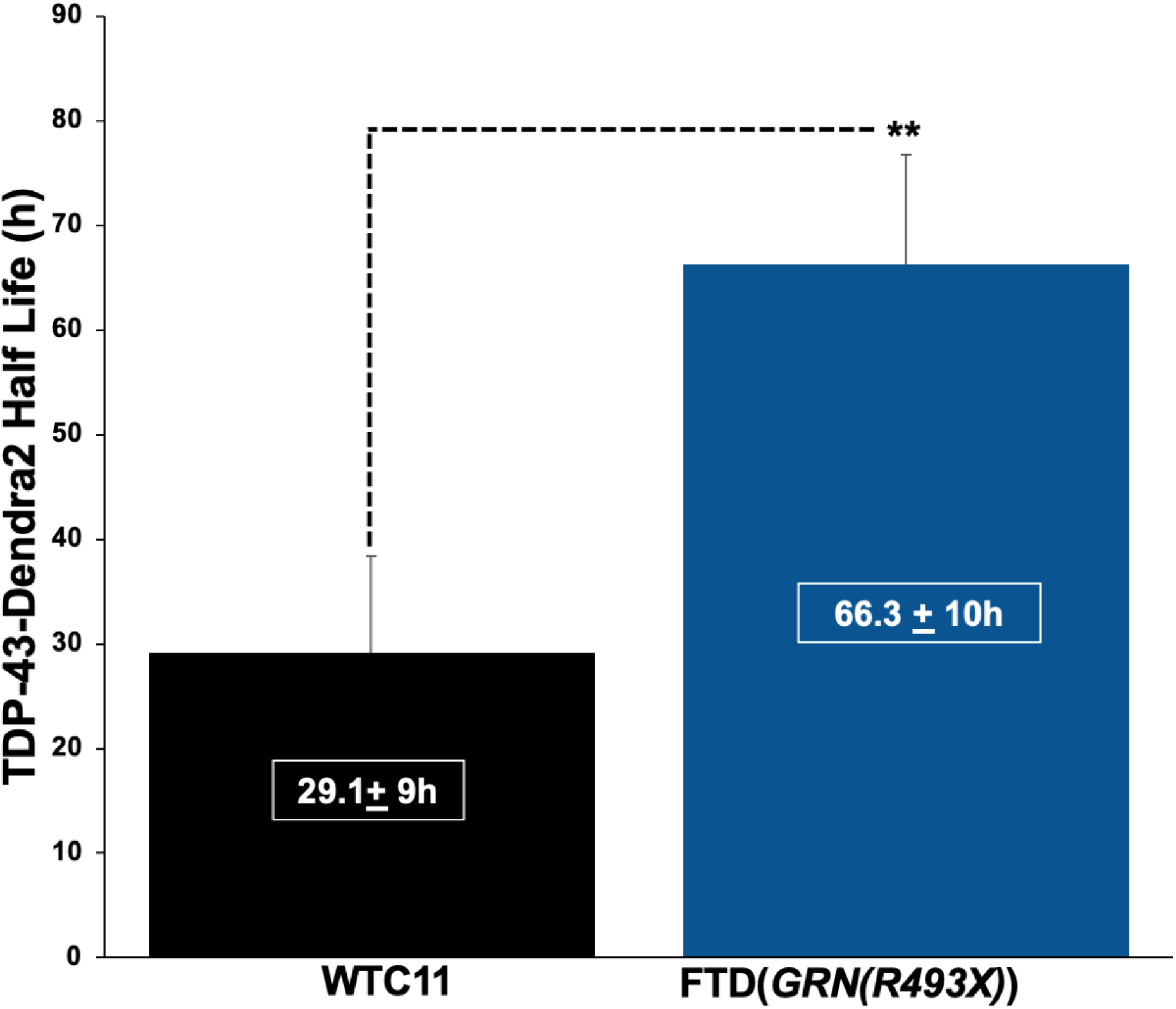
Clearance of TDP-43 is perturbed in PGRN-deficient FTD patient i-neurons. Measurement of TDP-43 turnover using the TDP-43-Dendra2 photoswitchable probe in human FTD (*GRN(R493X)*) patient or WTC11 healthy control cortical i-neurons imaged every 4h for 24h by longitudinal live cell RM. Statistical test: Student’s t-test (**p<0.01).

*GRN* has been identified in a GWAS study as a risk factor for PD, and PD is characterized by abnormal α-synuclein accumulation, suggesting that PGRN-deficiency may have a broader impact on the clearance of disease-causing proteins than its effects on TDP-43. To examine if PGRN-deficiency-induced protein dyshomeostasis may be a common thread in neurodegenerative disease, we also used an OPL approach to examine the turnover of α-synuclein in PGRN-deficient FTD i-neurons (**Supplemental Fig. 1**)^33^. Here we leveraged another photoswitchable protein, EOS3.2, which behaves similarly to Dendra2, fused to the human α-synuclein protein (EOS3.2-α-synuclein). Healthy control i-neurons had an average EOS3.2-α-synuclein half-life of ∼32h, whereas the PGRN-deficient FTD i-neurons had an average EOS3.2-α-synuclein half-life of ∼57h, indicating that α-synuclein turnover is impaired in the FTD i-neurons

### Lysosomes are enlarged and under-acidified in PGRN-deficient FTD i-neurons

Having discovered that PGRN-deficient FTD i-neurons exhibit phenotypes of neurodegeneration similar to those seen in patient brains, and had dysfunctional clearance of two known disease-associated proteins, TDP-43 and α-synuclein, we next investigated mechanisms by which *GRN* haploinsufficiency may lead to impaired protein clearance and neurodegeneration. We employed immunocytochemistry to test whether lysosomes were abnormal in the PGRN-deficient FTD i-neurons (**Fig. 5**). PGRN-deficiency is thought to compromise lysosome function and result in the accumulation of partially digested macromolecules and lipofuscin, leading to enlarged lysosomes. We found that the PGRN-deficient FTD i-neurons had an increase in enlarged Lamp1^+^ lysosomes (**Fig.5 E-H, I**) compared to healthy control i-neurons (**Fig. 5 A-D, I**). Interestingly, lysosomes were clustered in the soma of the PGRN-deficient FTD i-neurons in contrast to healthy control i-neurons, in which the lysosomes were more uniformly dispersed intracellularly.

**Figure 5:**
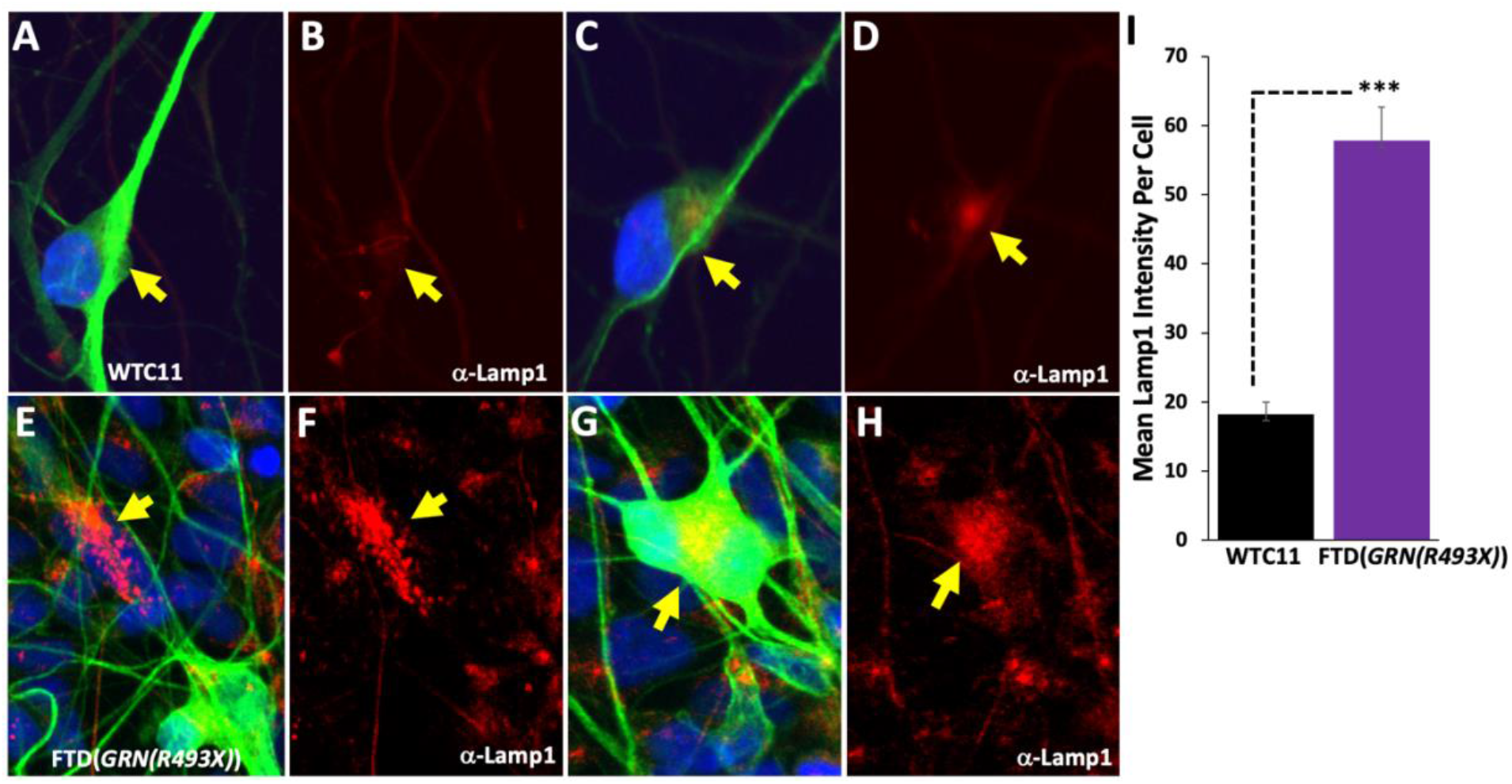
PGRN deficiency leads to enlarged lysosomes in human PGRN-deficient FTD patient i-neurons. Images of cultured human cortical i-neurons transduced with a lentivirus expressing EGFP (green) under the human synapsin1 promoter fixed and stained with DAPI (blue) and a Lamp1 antibody (red). **(A-D)**: Representative images of healthy control (WTC11) cortical neurons. **(E-H)**: Representative images of FTD patient (*GRN(R493X)*) cortical i-neurons. Yellow arrows indicate cell soma-localized lysosomes. **(I)** Quantitation of mean Lamp1 intensity per cell. Statistical test: Student’s t-test (***p<0.001).

PGRN is localized to and may provide functional support for lysosomes at several key points, including trafficking of degradative enzymes to lysosomes, and possibly regulating lysosomal acidification, as reported by Tanaka et al. (2017)^34^. Lysosomal acidification is important for the activation of degradative enzymes within this organelle. Deficits in these functions can lead to inefficient digestion and accumulation of cargo delivered to lysosomes for degradation, and result in lysosome enlargement as we found in the FTD i-neurons. Defective lysosomal acidification is a consistent feature underlying the pathobiology of lysosomal storage disorders (LSDs), including NCLs^35^.

To begin to understand mechanisms impacting lysosome function that may be disrupted by reduced PGRN levels, we measured the pH of lysosomes (pHlys) in the PGRN-deficient FTD i-neurons using the pHLARE biosensor. (**Table 1**)^36^. pHLARE is a genetically encoded ratiometric pHlys biosensor, that encodes LAMP1 tagged at its luminal amino-terminus with superfolder GFP (sfGFP) a GFP variant having a pKa ∼5.9, and tagged at the cytoplasmic carboxyl-terminus with mCherry. pHLARE is localized to lysosomes, and responds to pH changes in the lysosomal compartment with a dynamic range from pH 4.0 to 7.0. We first generated a pHlys calibration curve by expressing pHLARE in healthy control i-neurons treated with the protonophore, nigericin, and incubated the cells in buffers ranging from pH 4.0 up to pH 7.0, using RM to collect sfGFP and mCherry fluorescence signals. The fluorescence ratios from cells treated at each known pH value were calculated and then calibrated to absolute pHlys. We next used pHLARE to determine the average pHlys from healthy control cells and found that they exhibited an acidic pHlys of 4.5+0.2 (**Table 1**), which is comparable to that reported by Webb et al. (2021)^36^. A pHlys of ∼4.0-5.0 is critical for the optimal activity of lysosomal degradative enzymes. To further validate pHLARE as a pH sensor in live cells, we tested whether it accurately reported the neutralization of lysosomal luminal pH by bafilomycin A1 (BafA1), which is a specific inhibitor of the H+ ATPase (V-ATPase) that is required to acidify lysosomes. Treatment of the healthy control i-neurons expressing pHLARE with 50nM BafA1 increased the pHlys to 6.8+0.6, indicative of lysosomal under-acidification. Finally, we used pHLARE to measure the pHlys in PGRN-deficient FTD i-neurons and found they had an elevated pHlys of 6.8+0.7, suggesting that reduced PGRN leads to under-acidification of lysosomes in the PGRN-deficient FTD i-neurons, which is expected to impair the activity of multiple lysosomal hydrolases.

**TABLE 1.** Lysosomal pH Is Elevated In Human PGRN Deficient FTD Patient i-Neurons Cell Line Lysosomal pH p-value.

| **Cell Line** | **Lysosomal pH** | **p-value** |
| --- | --- | --- |
| Healthy Control (WTC11) i-Neurons, DMSO | 4.5±0.2 | ----- |
| Healthy Control (WTC11) i-Neurons, BafA1 (50nM) | 6.8±0.6 | *** |
| FTD Patient (GRN(R493X)), DMSO | 6.8±0.7 | *** |

Lysosomal pH reported as the mean ± s.d. Statistical test: Student's t-test (\*\*p<0.001).

### Quantitative TMT-proteomics reveal changes in protein homeostasis pathways in PGRN-deficient FTD i-neurons

Having shown that PGRN-deficient FTD i-neurons exhibit impaired protein clearance indicated by deficits in the turnover of both TDP-43 and α-synuclein, and the display aberrantly enlarged lysosomes, we wanted to determine if PGRN deficiency may impact the global neuronal proteome. One might expect, given FTD i-neurons exhibit lysosome dysfunction, that the metabolism of many proteins in these cells could be affected. To study the proteome of the FTD i-neurons we employed Tandem mass tag (TMT)-based quantitative global proteomics, a powerful, unbiased, and high-throughput approach to identify changes in protein expression that can provide insight into disease-related mechanisms associated with PGRN deficiency. We performed TMT-proteomics in the human PGRN-deficient FTD i-neurons (**Fig. 6**). A total of 7739 proteins were quantified. The FTD patient i-neurons had 324 proteins that were significantly decreased and 611 proteins that were significantly elevated. Gene Ontology (GO) analysis of all proteins whose levels were significantly altered in the PGRN-deficient FTD i-neurons were compared to control neurons using clusterProfiler. GO analysis of the upregulated proteins in the PGRN-deficient FTD i-neurons identified a significant enrichment of proteins associated with mitochondrial and endoplasmic reticulum (ER) terms. GO analysis for the downregulated proteins in the PGRN-deficient FTD i-neurons identified proteins associated with lipid, glycerolipid, and glycerophospholipid catabolism, lysosomal function, endosomes, cargo adaptor activity, vesicle tethering and transport, and in proteasome-mediated, ubiquitin-dependent protein catabolism, suggesting underlying perturbation of lipid and protein clearance pathways, including the ALP and UPS.

**Figure 6:**
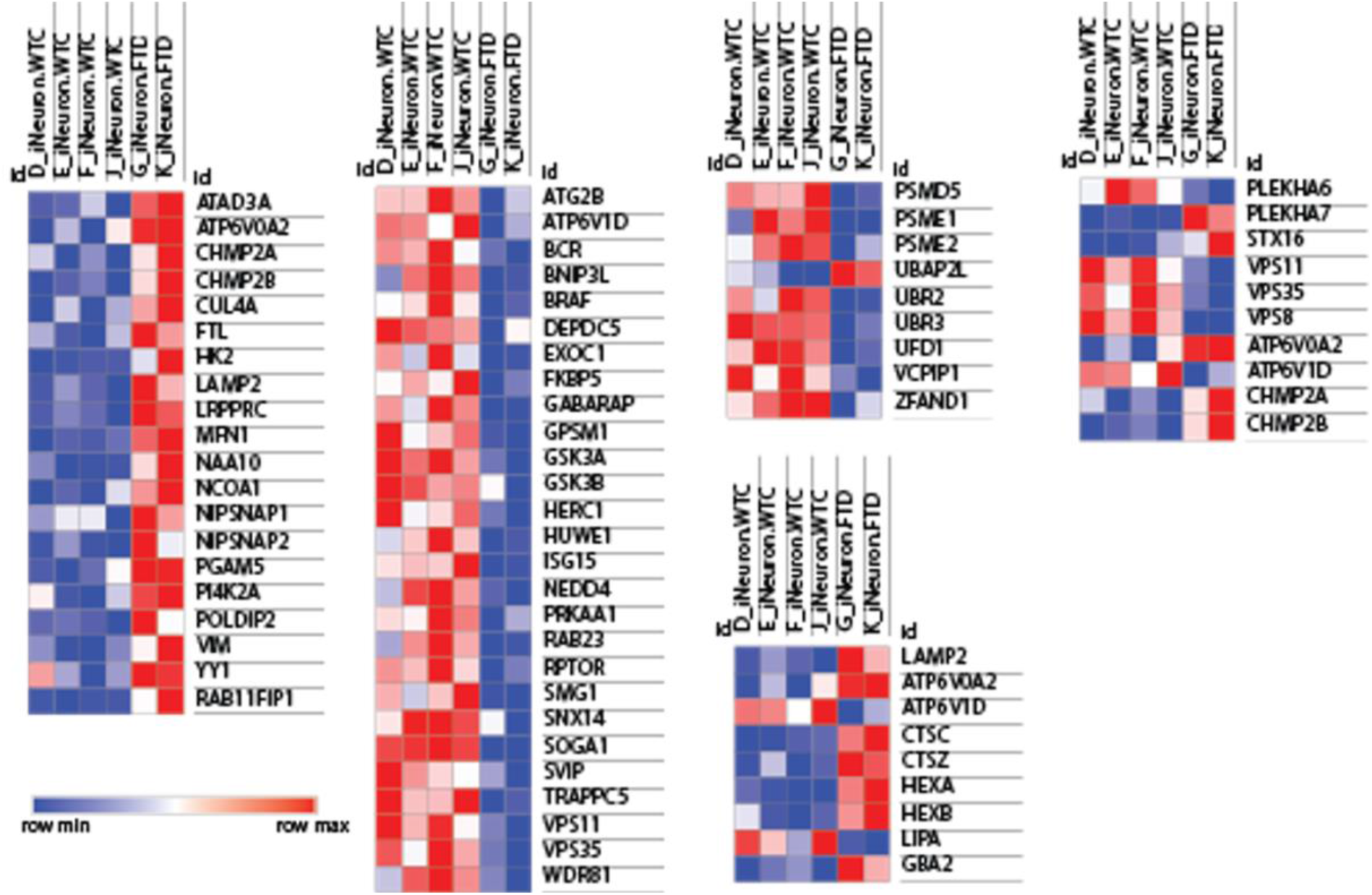
TMT-quantitative proteomics reveal changes in components of the autophagy-lysosome and proteasome clearance pathways in PGRN-deficient FTD patient i-neurons. TMT-quantitative proteomics analysis was performed on control (WTC11) or FTD (*GRN(R493X)*) patient cortical i-neurons with a total of 7739 total proteins quantified. Of those proteins, changes in the levels of 940 proteins were detected in the FTD patient i-neurons (p values <0.05). Results shown of 3 independent studies on a control line (WTC11) from a healthy individual and 2 independent studies on a FTD patient (*GRN(R493X)*) line.

Closer analysis of individual proteins showed significant downregulation of over 48 proteins linked to the ALP in the PGRN-deficient FTD i-neurons (**Fig. 6**), including a subset of CLEAR network proteins. We found that PGRN-deficient FTD i-neurons had significantly altered levels of 8 CLEAR network proteins compared to healthy control neurons including HEXA and HEXB (both increased), LAMP2 (increased), ATP6V1D (decreased), GABARAP (decreased), VPS8 (decreased), VPS11 (decreased), and the retromer subunit VSP35 (decreased). ATP6V1D is the catalytic domain subunit of the vATPase required for lysosome acidification. A second vATPase subunit, ATP6V0A2, was significantly increased in the PGRN-deficient FTD i-neurons (**Table 1**).

We also found changes in a small subset of lysosomal enzymes involved in protein degradation. Two lysosomal cysteine proteases, cathepsin C (CTSC) and cathepsin Z (CTSZ), were both increased in PGRN-deficient FTD i-neurons. In addition, we found that an endogenous inhibitor of cysteine proteases, cystatin C (CST3), was also increased in the PGRN-deficient FTD i-neurons.

We also identified changes in enzymes involved in lipid metabolism. Lysosomal acid lipase (LIPA), which is involved in the hydrolysis of cholesterol esters and triglycerides to release cholesterol and fatty acids, was reduced in the PGRN-deficient FTD i-neurons. We also found a set of enzymes involved in ganglioside metabolism were altered in the PGRN-deficient FTD i-neurons. Glucosylceramidase beta 2 (GBA2) was increased in the PGRN-deficient FTD i-neurons. GBA2 is a non-lysosomal glucosylceramidase related to the lysosomal glucosylceramidase GBA1 (GCase), which is linked to Gaucher’s disease. GBA1 levels were unchanged in the PGRN-deficient FTD i-neurons, suggesting that the changes in GBA2 in the FTD i-neurons were selective. Two other lysosomal enzymes important for ganglioside metabolism, hexaminidase A (HEXA) and hexaminidase B (HEXB), were both elevated in the PGRN-deficient FTD i-neurons

We also found that the expression of a subset of proteins linked to the proteasome pathway were altered in the PGRN-deficient FTD i-neurons. Two proteasome activator subunits, proteasome activator complex subunit 1 (PSME1) and proteasome activator complex subunit 2 (PSME2), were both reduced in the PGRN-deficient FTD i-neurons. The 26S Proteasome Non-ATPase Regulatory Subunit 5 (PSMD5) was also reduced in the PGRN-deficient FTD i-neurons. In addition, we found reduced levels of two E3 ubiquitin ligases, UBR2 and UBR3, in the PGRN-deficient FTD i-neurons. Changes in the levels of two proteins linked to the valosin-containing protein (VCP) were found in the PGRN-deficient FTD i-neurons: Valosin Containing Protein Interacting Protein 1 (VCPIP1), a deubiquitinating enzyme (DUB), and Ubiquitin Recognition Factor In ER-Associated Degradation Protein 1 (UFD1) were both reduced. Finally, we found reciprocal changes in two other proteasome pathway-linked proteins that have key roles in stress granule (SG) formation and turnover: ubiquitin-associated protein 2 (UBAP2L) was increased, and Zinc Finger AN1-Type Containing 1 (ZFAND1) was decreased in PGRN-deficient FTD i-neurons. These data raise the possibility that the two arms of protein clearance, the proteasome and the ALP may be impaired in PGRN-deficient FTD.

## Discussion

We have developed a human PGRN-deficient FTD patient neuronal cell model that replicates key aspects of the neuropathology associated with the disease. The i-neurons differentiated from FTD patient iPSCs have reduced PGRN levels, exhibit defective turnover and cytoplasmic accumulation of TDP-43, the most prominent ubiquitinated protein found in intracellular inclusions associated with FTD, as well as defective turnover of the PD-associated protein, α-synuclein, and display lysosome deficits. In addition, the human FTD cell models exhibit features of neurite loss and increased probability of death, indicating that the cells undergo spontaneous neurodegeneration.

The defective clearance of TDP-43 and α-synuclein is consistent with the PGRN-deficiency-linked lysosome dysfunction apparent in this cell model. Misfolded, aggregated TDP-43 or α-synuclein can be cleared by the lysosome, although both proteins can also be cleared by the proteasome^18-21, 37, 38^. Accumulation of either TDP-43 or α-synuclein, in turn, can negatively impact protein clearance pathways^39-41^. Proteomics performed in the FTD patient-derived i-neurons revealed altered levels of proteins linked to the ALP. Lysosomal dysfunction was implicated by increases in the levels of two lysosomal cysteine cathepsin proteases, CTSC and CTSZ along with cystatin C in the FTD patient i-neurons. Increased levels of lysosomal enzymes may signify compensation for inefficient degradation. The FTD patient i-neurons have impaired protein degradation despite elevated levels of cathepsin enzymes, suggesting that either cathepsin activation is compromised or compensatory mechanisms are not sufficiently robust to restore hydrolytic function. Cathepsin activation can be blocked by direct inhibition with a specific inhibitor or by disruption of its maturation. We found that cystatin C is elevated in the FTD patient i-neurons. Cystatin C is an endogenous cysteine protease inhibitor. This raises the possibility that this endogenous inhibitor may contribute to dysfunctional cathepsin activity. Alternatively, cathepsins are translated as proteolytically inactive enzymes containing inhibitory propeptides that are proteolytically removed to form a mature, active enzyme^42, 43^. Cathepsin maturation is often carried out by other cathepsins and their activity requires an acidified environment to proceed. We found altered levels of two subunits of the lysosomal v-ATPase, indicating that the lysosomal acidification machinery may be dysfunctional in FTD i-neurons. Accordingly, our pHLys determinations showing the PGRN-deficient FTD i-neurons expressed alkalized lysosomes, which would be expected to reduce the proteolytic capacity of the organelle.

The proteomics analysis also revealed that other arms of the proteostasis network may be aberrantly impacted or dysfunctional in the FTD patient i-neurons. We found the levels of LAMP2 were elevated in the FTD patient i-neurons, suggesting that CMA may be dysregulated. We also found a small subset of proteins linked to the UPS were altered in the PGRN-deficient FTD patient i-neurons. Both PSME1 and PSME2, which form a proteasomal activator complex, were reduced. Interestingly, we found reduced levels of two proteins linked to VCP–VCPIP1 and UFD1. VCP facilitates the degradation of polyubiquitylated substrates in the proteasome, and has known links to FTD^44^. We also found reciprocal changes in two other UPS-linked proteins that have key roles in stress granule (SG) formation and turnover, UBAP2L and ZFAND1. SGs are important in FTD because they have been implicated in TDP-43 mislocalization. UBAP2L is a ubiquitin binding protein that associates with SGs. In fact, UBAP2L has been shown to nucleate SGs in the presence of arsenite, endoplasmic reticulum, or heat stress^45^. ZFAND1 also plays a role in regulating SG turnover through the recruitment of VCP and the 26S proteasome^46^. Together, these data raise the possibility that both major arms of protein clearance, the ALP and the UPS, may be impaired in PGRN-deficient FTD. It remains to be determined whether the lysosomal deficits lead to dysregulation of the UPS, or if the UPS is independently affected by *GRN*-haploinsufficiency. Future studies will explore what are the interrelationships between proteasome and lysosome clearance pathways and what are the underlying mechanisms contributing to proteostasis impairment in PGRN-deficient FTD patient i-neurons.

Proteomics also identified dysregulation of glycerolipid catabolism in the FTD patient i-neurons, and our results are consistent with other proteomics studies using mouse models of PGRN-deficiency. Both Huang et al. (2020)^47^ and Boland et al. (2021)^48^ found changes in proteins associated with lipid catabolism in *Grn*-deficient mouse brain samples^47, 48^. We found reduced levels of lysosomal acid lipase (LIPA), which is involved in the hydrolysis of cholesterol esters and triglycerides to release cholesterol and fatty acids, in the PGRN-deficient FTD i-neurons. We also found that changes in proteins involved in metabolizing gangliosides correlated with PGRN-deficiency. GBA2, a non-lysosomal glucosylceramidase related to the lysosomal glucosylceramidase GBA (GCase), was increased in the PGRN-deficient FTD i-neurons. PGRN-deficiency has been shown to perturb GCase activity, however GBA1 levels were unchanged in the PGRN-deficient FTD i-neurons, suggesting that the changes in GBA2 levels were selective for the FTD i-neurons. Interestingly, in a mouse model of Neimann Pick Type C disease (NPC), reducing GBA2 levels was beneficial to neurons, suggesting that the increase in GBA2 levels in the FTD patient i-neurons may have deleterious effects^49^.

We also found both HEXA and HEXB were increased in the PGRN-deficient FTD patient i-neurons. These enzymes are both involved in ganglioside metabolism. PGRN has been reported to bind directly to HEXA^50^. In PGRN-deficient cells, HEXA has been shown to aggregate and its substrate, GM2 ganglioside, accumulates, leading to an LSD phenotype. Future studies will focus on how PGRN-deficiency results in aberrant lipid metabolism, and whether disruption of lipid homeostasis negatively impacts protein trafficking and/or turnover and further contribute to the neuropathology of FTD.

## Materials and Methods

### Reagents and antibodie

pGW1-TDP-43-Dendra2 is described in Barmada et al. (2014)^31^. pGW1-EOS3.2-a-synuclein was derived from pGW1-Dendra2-a-synuclein described in Skibinski et al. (2016)^33^ where Dendra2 was replaced the photoswitchable protein, EOS3.2^33, 51^. The rabbit anti-TDP-43 polyclonal antibody is from Proteintech (10782-2-AP; 1:1000). The rabbit anti-Lamp1 polyclonal antibody is from abcam (ab24170; 1:300).

### Cortical forebrain i-neuron differentiation from iPSC

Cortical i-neurons were differentiated from human FTD patient iPSCs harboring the GRN(R493X) mutation using a modification of the method reported in Boissart et al. (2013)^52^. iPSCs were cultured to confluency as a monolayer then switched into medium containing LDN193189 and SB431542 to induce dual SMAD inhibition and commit iPSCs to a neural lineage. The differentiation medium was changed every 24h until day 11. At day 11, neural rosettes containing neural precursor cells (NPCs) were formed. The NPCs were passaged into N2B27 medium containing epidermal growth factor (EGF; 10ng ml^-1^), fibroblast growth factor-2 (FGF2; 10 ng ml^-1^) and brain-derived growth factor (BDNF; 20 ng ml^-1^). Confluent cells were passed onto growth factor reduced Matrigel using Accutase first as small multicellular clumps (passages 2–4) at a ratio of 1:3, then as a single-cell suspension with a density of 100,000 cells cm^-2^.

Neuronal differentiation of the NPCs was induced by plating the cells at low density (50,000 cells cm^-2^) onto poly-ornithine/laminin/fibronectin-treated multiwell culture plates, in N2B27 supplemented with BDNF (20ng ml^-1^) and glial cell derived neurotrophic factor (GDNF; 20 ng ml^-1^) and without EGF and FGF. The medium was changed every 4 days.

### OPL turnover of TDP-43 or α-synuclein

Turnover of either transfected pGW1-TDP-43-Dendra2 or pGW1-EOS3.2-α-synuclein was performed by optical pulse labeling (OPL). Dendra2 and EOS3.2 are photoswitchable proteins. The Dendra2 or EOS3.2 proteins were photoswitched using a 4s pulse of 405nm light, then cells were imaged longitudinally every 4h using robotic microscopy over 24h. Green and red fluorescence images were captured. The linear decay of the red fluorescent photoswitched population of molecules was used to calculate the mean half-life of either TDP-43-Dendra2 or EOS3.2-α-synuclein.

### pHLARE^36^

pGW1-pHLARE was transfected into 36 DIV early mature cortical i-neurons, and subjected to longitudinal live cell imaging by robotic microscopy at 40 DIV. In order to calculate fluorescence ratios of pHLARE, we followed the approach reported in Webb et al. (2021)^36^. Briefly, cells were incubated in nigericin buffers (pH 4.0-pH 8.0) for 5 min before imaging to equilibrate extracellular and pHi. Nigericin incubations to calibrate fluorescence ratios to pHlys were included to obtain a calibration conversion formula for each imaging experiment and for each cell.

### Quantitative Tandem Mass Tag (TMT) proteomics

Flash frozen NPC and i-neuron cell pellets were lysed in 8M urea buffer (8M urea, 150 mM NaCl, 50 mM HEPES pH 7.5, 1x EDTA-free protease inhibitor cocktail [Roche], 1x PhosSTOP phosphatase inhibitor cocktail [Roche]). Lysates were clarified by centrifugation at 17,000 x g for 15 min at 4°C. Protein concentration of the supernatant was quantified by bicinchroninic acid assay ([BCA], Pierce). To reduce and alkylate cysteines, 150 μg of protein was sequentially incubated with 5 mM TCEP for 30 mins, 14 mM iodoacetamide for 30 mins, and 10 mM DTT for 15 mins. All reactions were performed at RT. Next, proteins were chloroform-methanol precipitated and the pellet resuspended in 200 mM EPPS pH 8.5. Then, the protease LysC (Wako) was added at 1:100 (LysC:protein) ratio and incubated overnight at RT. The day after, samples were further digested for 5 hours at 37ºC with trypsin at 1:75 (trypsin:protein) ratio. Both digestions were performed in an orbital shaker at 1,500 rpm. After digestion, samples were clarified by centrifugation at 17,000 x g for 10 min. Peptide concentration of the supernatant was quantified using a quantitative colorimetric peptide assay (Thermo Fisher). For TMT labelling we used the TMT-11plex kit (Thermo Fisher)^53, 54^. Briefly, 25 μg of peptides was brought to 1 μg/μl with 200 mM EPPS (pH 8.5), acetonitrile (ACN) was added to a final concentration of 30% followed by the addition of 50 μg of each TMT reagents. After 1 h of incubation at RT, the reaction was stopped by the addition of 0.3% hydroxylamine (Sigma) for 15 min at RT. After labelling, samples were combined, desalted with tC18 SepPak solid-phase extraction cartridges (Waters), and dried in the SpeedVac. Next, desalted peptides were resuspended in 5% ACN, 10 mM NH_4_HCO_3_ pH 8 and fractionated in a basic pH reversed phase chromatography using a HPLC equipped with a 3.5 μm Zorbax 300 Extended-C18 column (Agilent). Fractions were collected in a 96-well plate, then combined into 24 samples. Twelve of them were desalted following the C18 Stop and Go Extraction Tip (STAGE-Tip) ^55^ and dried down in a SpeedVac. Finally, peptides were resuspended in 1% formic acid, 3% ACN, and analyzed by LC-MS3 in an Orbitrap Fusion Lumos (Thermo Fisher) mounted with FAIMS and running in high-resolution MS2 (hrMS2) mode^56, 57^. The scan sequence began a MS1 spectrum (Orbitrap analysis, resolution 60,000, 400–1600 Th, 400,000 automatic gain control (AGC) target, maximum injection time 50 ms) followed by MS2 analysis consisted of higher-energy collisional dissociation (HCD), 125,000 AGC target, NCE (normalized collision energy) was 37%, resolution was 50,000, maximum injection time was 150ms, and isolation window was 0.5 Th. FAIMS CVs were set to −40, −60, and −80 V.

### Mass spectrometry data analysis

A suite of in-house pipeline (GFY-Core Version 3.8, Harvard University) was used to obtain final protein quantifications from all RAW files collected. RAW data were converted to mzXML format using MSconvert^58^ and searched using the search engine Comet^59^ against a human target-decoy protein database (downloaded from UniProt in June 2019) that included the most common contaminants. Precursor ion tolerance was set at 20 ppm and product ion tolerance at 0.02 Da. Cysteine carbamidomethylation (+57.0215 Da) and TMT tag (+229.1629 Da) on lysine residues and peptide N-termini were set as static modifications. Up to 2 variable methionine oxidations (+15.9949 Da) and 2 miss cleavages were allowed in the searches. Peptide-spectrum matches (PSMs) were adjusted to a 1% FDR with a linear discriminant analysis^60^ and proteins were further collapsed to a final protein-level FDR of 1%. TMT quantitative values we obtained from MS2 scans. Only those with a signal-to-noise ratio > 100 and an isolation specificity > 0.7 were used for quantification. Each TMT was normalized to the total signal in each column. Quantifications are represented as relative abundances. RAW files will be made available upon request.

### Experimental design and statistical analysi

Cumulative risk of death curves compiled from ten separate experiments, including a total of 38,382 FTD*(GRN(R493X)*) patient i-neurons and 36,106 healthy control WTC11 i-neurons expressing EGFP under the human synapsin1 promoter.

All other experiments were performed at least two times. Statistical analysis was performed using Excel. Data from two conditions were compared using two-tailed Student’s t-test. The level of significance was set at < 0.05. Quantitative data are presented as mean + s.d.

## Acknowledgments

We thank Rick Morimoto, Judith Frydman, Jeff Kelly, Evan Powers, Dan Garza and members of the Proteostasis Consortium, and members of the Finkbeiner lab for discussion and comments on these experiments. We thank Terry Reisine for comments and editorial assistance, and Gayane Abramova and Kelley Nelson for administrative assistance. Study funding: Primary support for this work was from P01AG054407 and RF1AG060765. Additional support came from RF1NS128800 and R01LM013617, the Taube/Koret Center for Neurodegenerative Disease Research, and the J. David Gladstone Institutes. M.A.P. was supported by the Goldberg Fund Fellowship and the Miguel Servet Program (CP21/00017).

## References

1. Chen Y, Li S, Su L, Sheng J, Lv W, Chen G, Xu Z. (2015) Association of progranulin polymorphism rs5848 with neurodegenerative diseases: A meta-analysis. J. Neurol. 262:814–822.

2. Xu HM, Tan L, Wan Y, Tan MS, Zhang W, Zheng ZJ, Kong LL, Wang ZX, Jiang T, Tan L, Yu JT. (2017) PGRN is associated with late-onset Alzheimer’s disease: A case-control replication study and meta-analysis. Mol. Neurobiol. 54:1187–1195.

3. Perry DC, Lehmann M, Yokoyama JS, Karydas A, et al. (2013) Progranulin mutations as risk factors for Alzheimer disease. JAMA Neurol. 70:774–778. PMCID: PMC3743672

4. Hosokawa M, Arai T, Masuda-Suzukake M, Kondo H, Matsuwaki T, Nishihara M, Hasegawa M, Akiyama H. (2015) Progranulin reduction is associated with increased tau phosphorylation in P301L tau transgenic mice. J. Neuropathol. Exp. Neurol. 74:158–165.

5. Hsiung GY, Fok A, Feldman HH, Rademakers R, Mackenzie IR. (2011) rs5848 polymorphism and serum progranulin level. J. Neurol. Sci. 300:28–32. PMCID: PMC3085023

6. Kamalainen A, Viswanathan J, Natunen T, Helisalmi S, et al. (2013) GRN variant rs5848 reduces plasma and brain levels of granulin in Alzheimer’s disease patients. J. Alzheimers Dis. 33:23–27.

7. Minami SS, Min SW, Krabbe G, Wang C, et al. (2014) Progranulin protects against amyloid β deposition and toxicity in Alzheimer’s disease mouse models. Nat. Med. 20:1157–1164. PMCID: PMC4196723

8. Pereson S, Wils H, Kleinberger G, McGowan E, et al. (2009) Progranulin expression correlates with dense-core amyloid plaque burden in Alzheimer disease mouse models. J. Pathol. 219:173–181.

9. Nalls MA, Blauwendraat C, Vallerga CL, Heilbron K, et al. (2019) Identification of novel risk loci, causal insights, and heritable risk for Parkinson’s disease: A meta-analysis of genome-wide association studies. Lancet. Neurol. 18:1091–1102.

10. Leverenz JB, Yu CE, Montine TJ, Steinbart E, Bekris LM, Zabetian C, Kwong LK, Lee VM, Schellenberg GD, Bird TD. (2007) A novel progranulin mutation associated with variable clinical presentation and tau, TDP43 and alpha-synuclein pathology. Brain 130:1360–1374.

11. Smith KR, Damiano J, Franceschetti S, Carpenter S, et al. (2012) Strikingly different clinicopathological phenotypes determined by progranulin-mutation dosage. Am. J. Hum. Genet. 90:1102–1107. PMCID: PMC3370276

12. Faber I, Prota JR, Martinez AR, Lopes-Cendes I, França MCJ. (2017) A new phenotype associated with homozygous GRN mutations: Complicated spastic paraplegia. Eur. J. Neurol. 24:e3–e4.

13. Kamate M, Detroja M, Hattiholi V. (2019) Neuronal ceroid lipofuscinosis type-11 in an adolescent. Brain Dev. 41:542–545.

14. Arrant AE, Onyilo VC, Unger DE, Roberson ED. (2018) Progranulin gene therapy improves lysosomal dysfunction and microglial pathology associated with frontotemporal dementia and neuronal ceroid lipofuscinosis. J. Neurosci. 38:2341–2358. PMCID: PMC5830520

15. Jian J, Zhao S, Tian QY, Liu H, et al. (2016) Association between progranulin and Gaucher disease. EBioMedicine 11:127–137. PMCID: PMC5049935

16. Jian J, Tian QY, Hettinghouse A, Zhao S, et al. (2016) Progranulin recruits HSP70 to β-glucocerebrosidase and is therapeutic against Gaucher disease. EBioMedicine 13:212–224. PMCID: PMC5264254

17. Elia LP, Mason AR, Alijagic A, Finkbeiner S. (2019) Genetic regulation of neuronal progranulin reveals a critical role for the autophagy-lysosome pathway. J. Neurosci. 39:3332–3344. PMCID: PMC6788815

18. Ormeño F, Hormazabal J, Moreno J, Riquelme F, Rios J, Criollo A, Albornoz A, Alfaro IE, Budini M. (2020) Chaperone mediated autophagy degrades TDP-43 protein and is affected by TDP-43 aggregation. Front. Mol. Neurosci. 13:19. PMCID: PMC7040037

19. Scotter EL, Vance C, Nishimura AL, Lee YB, Chen HJ, Urwin H, Sardone V, Mitchell JC, Rogelj B, Rubinsztein DC, Shaw CE. (2014) Differential roles of the ubiquitin proteasome system and autophagy in the clearance of soluble and aggregated TDP-43 species. J. Cell Sci. 127:1263–1278. PMCID: PMC3953816

20. Cicardi ME, Cristofani R, Rusmini P, Meroni M, et al. (2018) TDP-25 routing to autophagy and proteasome ameliorates its aggregation in amyotrophic lateral sclerosis target cells. Sci. Rep. 8:12390. PMCID: PMC6098007

21. Zhang YJ, Gendron TF, Xu YF, Ko LW, Yen SH, Petrucelli L. (2010) Phosphorylation regulates proteasomal-mediated degradation and solubility of TAR DNA binding protein-43 C-terminal fragments. Mol. Neurodegener. 5:33. PMCID: PMC2941488

22. Yin F, Banerjee R, Thomas B, Zhou P, et al. (2010) Exaggerated inflammation, impaired host defense, and neuropathology in progranulin-deficient mice. J. Exp. Med. 207:117–128. PMCID: PMC2812536

23. Martens LH, Zhang J, Barmada SJ, Zhou P, Kamiya S, Sun B, Min SW, Gan L, Finkbeiner S, Huang EJ, Farese RV, Jr. (2012) Progranulin deficiency promotes neuroinflammation and neuron loss following toxin-induced injury. J. Clin. Invest. 122:3955–3959. PMCID: PMC3484443

24. Kayasuga Y, Chiba S, Suzuki M, Kikusui T, Matsuwaki T, Yamanouchi K, Kotaki H, Horai R, Iwakura Y, Nishihara M. (2007) Alteration of behavioural phenotype in mice by targeted disruption of the progranulin gene. Behav. Brain Res. 185:110–118.

25. Petkau TL, Neal SJ, Milnerwood A, Mew A, et al. (2012) Synaptic dysfunction in progranulin-deficient mice. Neurobiol. Dis. 45:711–722.

26. Filiano AJ, Martens LH, Young AH, Warmus BA, et al. (2013) Dissociation of frontotemporal dementia-related deficits and neuroinflammation in progranulin haploinsufficient mice. J. Neurosci. 33:5352–5361. PMCID: PMC3740510

27. Arrant AE, Filiano AJ, Warmus BA, Hall AM, Roberson ED. (2016) Progranulin haploinsufficiency causes biphasic social dominance abnormalities in the tube test. Genes Brain Behav. 15:588–603. PMCID: PMC5943713

28. Yin F, Dumont M, Banerjee R, Ma Y, Li H, Lin MT, Beal MF, Nathan C, Thomas B, Ding A. (2010) Behavioral deficits and progressive neuropathology in progranulin-deficient mice: A mouse model of frontotemporal dementia. FASEB J. 24:4639–4647. PMCID: PMC2992364

29. Josephs KA, Ahmed Z, Katsuse O, Parisi JF, et al. (2007) Neuropathologic features of frontotemporal lobar degeneration with ubiquitin-positive inclusions with progranulin gene (PGRN) mutations. J. Neuropathol. Exp. Neurol. 66:142–151.

30. Arrasate M, Mitra S, Schweitzer ES, Segal MR, Finkbeiner S. (2004) Inclusion body formation reduces levels of mutant huntingtin and the risk of neuronal death. Nature 431:805–810.

31. Barmada SJ, Serio A, Arjun A, Bilican B, et al. (2014) Autophagy induction enhances TDP43 turnover and survival in neuronal ALS models. Nat. Chem. Biol. 10:677–685. PMCID: PMC4106236

32. Finkbeiner S, Frumkin M, Kassner PD. (2015) Cell-based screening: Extracting meaning from complex data. Neuron 86:160–174. PMCID: PMC4457442

33. Skibinski G, Hwang V, Ando DM, Daub A, Lee AK, Ravisankar A, Modan S, Finucane MM, Shaby BA, Finkbeiner S. (2017) Nrf2 mitigates LRRK2- and α-synuclein-induced neurodegeneration by modulating proteostasis. Proc. Natl. Acad. Sci. U. S. A. 114:1165–1170. PMCID: PMC5293055

34. Tanaka Y, Suzuki G, Matsuwaki T, Hosokawa M, Serrano G, Beach TG, Yamanouchi K, Hasegawa M, Nishihara M. (2017) Progranulin regulates lysosomal function and biogenesis through acidification of lysosomes. Hum. Mol. Genet. 26:969–988.

35. Bagh MB, Peng S, Chandra G, Zhang Z, Singh SP, Pattabiraman N, Liu A, Mukherjee AB. (2017) Misrouting of v-ATPase subunit V0a1 dysregulates lysosomal acidification in a neurodegenerative lysosomal storage disease model. Nat. Commun. 8:14612. PMCID: PMC5344305

36. Webb BA, Aloisio FM, Charafeddine RA, Cook J, Wittmann T, Barber DL. (2021) pHLARE: A new biosensor reveals decreased lysosome pH in cancer cells. Mol. Biol. Cell 32:131–142. PMCID: PMC8120692

37. Webb JL, Ravikumar B, Atkins J, Skepper JN, Rubinsztein DC. (2003) Alpha-synuclein is degraded by both autophagy and the proteasome. J. Biol. Chem. 278:25009–25013.

38. Mak SK, McCormack AL, Manning-Bog AB, Cuervo AM, Di Monte DA. (2010) Lysosomal degradation of alpha-synuclein in vivo. J Biol Chem 285:13621–13629. PMCID: PMC2859524

39. McKinnon C, De Snoo ML, Gondard E, Neudorfer C, et al. (2020) Early-onset impairment of the ubiquitin-proteasome system in dopaminergic neurons caused by α-synuclein. Acta Neuropathol. Commun. 8:17. PMCID: PMC7023783

40. Martinez-Vicente M, Talloczy Z, Kaushik S, Massey AC, et al. (2008) Dopamine-modified alpha-synuclein blocks chaperone-mediated autophagy. J. Clin. Invest. 118:777–788. PMCID: PMC2157565

41. Xia Q, Wang H, Hao Z, Fu C, Hu Q, Gao F, Ren H, Chen D, Han J, Ying Z, Wang G. (2016) TDP-43 loss of function increases TFEB activity and blocks autophagosome-lysosome fusion. EMBO j. 35:121–142. PMCID: PMC4718457

42. Lamort AS, Hamon Y, Czaplewski C, Gieldon A, et al. (2019) Processing and maturation of cathepsin C zymogen: A biochemical and molecular modeling analysis. Int. J. Mol. Sci. 20:PMCID: PMC6801622

43. Hamon Y, Legowska M, Hervé V, Dallet-Choisy S, et al. (2016) Neutrophilic cathepsin C is maturated by a multistep proteolytic process and secreted by activated cells during inflammatory lung diseases. J. Biol. Chem. 291:8486–8499. PMCID: PMC4861422

44. Wani A, Zhu J, Ulrich JD, Eteleeb A, et al. (2021) Neuronal VCP loss of function recapitulates FTLD-TDP pathology. Cell Rep. 36:109399. PMCID: PMC8383344

45. Cirillo L, Cieren A, Barbieri S, Khong A, Schwager F, Parker R, Gotta M. (2020) UBAP2L forms distinct cores that act in nucleating stress granules upstream of G3BP1. Curr. Biol. 30:698–707.e6.

46. Turakhiya A, Meyer SR, Marincola G, Böhm S, Vanselow JT, Schlosser A, Hofmann K, Buchberger A. (2018) ZFAND1 recruits p97 and the 26S proteasome to promote the clearance of arsenite-induced stress granules. Mol. Cell 70:906–919.e7.

47. Huang M, Modeste E, Dammer E, Merino P, et al. (2020) Network analysis of the progranulin-deficient mouse brain proteome reveals pathogenic mechanisms shared in human frontotemporal dementia caused by GRN mutations. Acta Neuropathol. Commun. 8:163. PMCID: PMC7541308

48. Boland S, Swarup S, Ambaw YA, Malia PC, et al. (2022) Deficiency of the frontotemporal dementia gene GRN results in gangliosidosis. Nat. Commun. 13:5924. PMCID: PMC9546883

49. Marques AR, Aten J, Ottenhoff R, van Roomen CP, et al. (2015) Reducing GBA2 activity ameliorates neuropathology in Niemann-Pick type C mice. PLoS One 10:e0135889. PMCID: PMC4537125

50. Chen Y, Jian J, Hettinghouse A, Zhao X, Setchell KDR, Sun Y, Liu CJ. (2018) Progranulin associates with hexosaminidase A and ameliorates GM2 ganglioside accumulation and lysosomal storage in Tay-Sachs disease. J. Mol. Med. (Berl) 96:1359–1373. PMCID: PMC6240367

51. Zhang M, Chang H, Zhang Y, Yu J, et al. (2012) Rational design of true monomeric and bright photoactivatable fluorescent proteins. Nat. Methods 9:727–729.

52. Boissart C, Poulet A, Georges P, Darville H, Julita E, Delorme R, Bourgeron T, Peschanski M, Benchoua A. (2013) Differentiation from human pluripotent stem cells of cortical neurons of the superficial layers amenable to psychiatric disease modeling and high-throughput drug screening. Transl. Psychiatry 3:e294. PMCID: PMC3756296

53. Cao C, Prado MA, Sun L, Rockowitz S, Sliz P, Paulo JA, Finley D, Fleming MD. (2021) Maternal iron deficiency modulates placental transcriptome and proteome in mid-gestation of mouse pregnancy. J. Nutr. 151:1073–1083. PMCID: PMC8112763

54. Nguyen AT, Prado MA, Schmidt PJ, Sendamarai AK, et al. (2017) UBE2O remodels the proteome during terminal erythroid differentiation. Science 357:eaan0218. PMCID: PMC5812729

55. Rappsilber J, Ishihama Y, Mann M. (2003) Stop and go extraction tips for matrix-assisted laser desorption/ionization, nanoelectrospray, and LC/MS sample pretreatment in proteomics. Anal. Chem. 75:663–670.

56. Schweppe DK, Rusin SF, Gygi SP, Paulo JA. (2020) Optimized workflow for multiplexed phosphorylation analysis of TMT-labeled peptides using high-field asymmetric waveform ion mobility spectrometry. J. Proteome Res. 19:554–560. PMCID: PMC6996458

57. Schweppe DK, Prasad S, Belford MW, Navarrete-Perea J, et al. (2019) Characterization and optimization of multiplexed quantitative analyses using high-field asymmetric-waveform ion mobility mass spectrometry. Anal. Chem. 91:4010–4016. PMCID: PMC6993951

58. Chambers MC, Maclean B, Burke R, Amodei D, et al. (2012) A cross-platform toolkit for mass spectrometry and proteomics. Nat. Biotechnol. 30:918–920. PMCID: PMC3471674

59. Eng JK, Jahan TA, Hoopmann MR. (2013) Comet: An open-source MS/MS sequence database search tool. Proteomics 13:22–24.

60. Huttlin EL, Jedrychowski MP, Elias JE, Goswami T, Rad R, Beausoleil SA, Villén J, Haas W, Sowa ME, Gygi SP. (2010) A tissue-specific atlas of mouse protein phosphorylation and expression. Cell 143:1174–1189. PMCID: PMC3035969

