## Supplemental Figure 1 for "Frontotemporal Dementia Patient Neurons With Progranulin Deficiency Display Protein Dyshomeostasis"

### Supplemental Figure Legend

**Supplemental Figure 1**: **Clearance of α-synuclein is perturbed in PGRN-deficient FTD patient i-neurons.** Measurement of α-synuclein turnover using the EOS3.2-α-synuclein photoswitchable probe in human FTD(*GRN(R493X)*) patient or WTC11 healthy control cortical i-neurons imaged every 4h for 24h by longitudinal live cell RM. Statistical test: Student's t-test (**p<0.01).

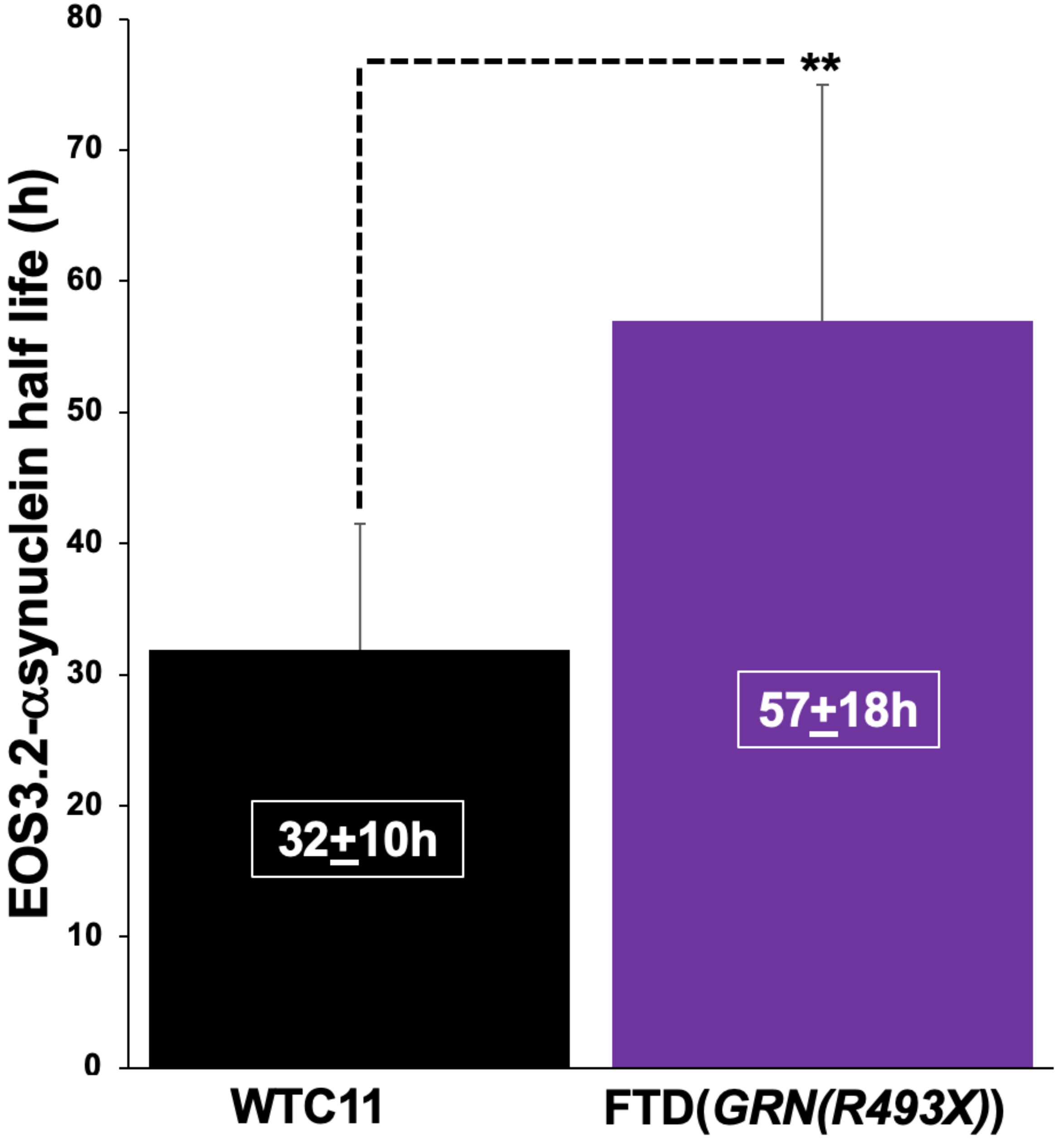


**Supplemental Figure 1**
